# Systematic Characterization of PBP2 as the Primary Siderophore Recognizer in Actinomycetes and Other Gram-Positive Bacteria

**DOI:** 10.1101/2025.09.29.679136

**Authors:** Linlong Yu, Guanyue Xiong, Zhiyuan Li

## Abstract

Iron is a scarce yet essential nutrient for bacteria. Microbes often acquire iron by secreting siderophores, a diverse group of small molecules that form high-affinity complexes with iron for microbial uptake. Understanding microbial iron interaction networks requires detailed characterization of siderophore recognition specificity. In Gram-positive bacteria, substrate-binding proteins (SBPs) bind iron-siderophore complexes and deliver them to ABC transporters for import. However, the SBPs responsible for selective recognition remain poorly characterized, hindering large-scale data mining and network reconstruction. Here, we addressed this knowledge gap by systematically analyzing siderophore uptake systems, first in five representative genera and then across a comprehensive dataset of 16,232 Gram-positive genomes. Through a pipeline integrating genome mining, coevolutionary analysis, and structural modeling, we established PBP2 (Peripla_BP_2) subtype SBPs as the primary siderophore recognizer family. We revealed that, unlike the physically clustered systems in Gram-negative bacteria, synthetase and recognizer genes in Gram-positive bacteria are sometimes genomically decoupled yet display coordinated transcriptional regulation by iron-responsive transcription factors. Our findings underscore key differences between Gram-positive and Gram-negative iron acquisition systems, providing foundational knowledge for large-scale inference of siderophore-mediated microbial interactions.

**Impact Statement:** Bacteria secrete siderophores to scavenge iron and import the siderophore-iron complex via specific receptors, a process that shapes microbial community dynamics. However, predicting these interactions has been challenging, because the specific siderophore receptors in Gram-positive bacteria remained largely uncharacterized. In this study, we opened this “black box” by analyzing a comprehensive dataset of 16,232 genomes spanning the majority of Gram-positive bacteria. Through coevolutionary analysis, we identified PBP2 proteins as the primary “locks” that recognize siderophore “keys.” We further demonstrate that these receptors exhibit greater evolutionary flexibility than their Gram-negative counterparts, frequently decoupled genomically from siderophore biosynthesis genes yet linked by transcriptional regulation. This discovery fills a critical knowledge gap, providing the missing link needed to map the global landscape of siderophore uptake potential and enable “sequence-to-ecology” prediction of iron-interaction networks in Gram-positive bacteria.

## Introduction

Iron is essential for bacterial growth, survival, and virulence[1], serving as a cofactor in key enzymes involved in DNA replication and electron transfer[2, 3]. Despite its abundance on Earth, iron’s bioavailability in natural environments is severely limited due to the oxidation of ferrous iron (Fe²⁺) to insoluble ferric iron (Fe³⁺) following the Great Oxidation Event [4–6]. To overcome this scarcity, most bacteria secrete siderophores to the surrounding environment to chelate iron[4]. Siderophores are a class of small, high-affinity iron-chelating secondary metabolites[1, 4, 7, 8], biosynthesized by non-ribosomal peptide synthetase (NRPS) or NRPS-independent siderophore synthetase (NIS) pathways[8–10]. Once secreted, siderophores scavenge iron from the environment to form siderophore-iron complexes, which are then imported back into bacterial cells through specific membrane recognizers[11–13].

The specificity between siderophores and the membrane proteins selectively recognizing them (i.e. recognizers) mediates intricate microbial interactions in iron competition[4]. Previous studies have shown that siderophores are preferentially bound and imported by specific receptors, akin to keys fitting locks[13–16]. For instance, three *Pseudomonas aeruginosa* strains produce chemically distinct pyoverdines and exhibit high specificity, primarily utilizing their own siderophores with minimal cross-recognition[13, 15]. Meanwhile, microbes often possess multiple membrane recognizers to pirate xenosiderophores produced by others. For example, the Gram-positive *Bacillus subtilis* produces Bacillibactin yet imports additional siderophores, including Enterobactin, Ferrichrome, and Ferrioxamine E[17–19]. This lock-and-key specificity likely evolves from an arms race between producers and cheaters, shaping rich community games: For microbes with matching receptors, a siderophore acts as a shared public good facilitating iron uptake; for those without, it becomes a “public bad” that sequesters iron away[20]. Understanding siderophore-recognizer specificity is key to dissecting complex microbial communities[4].

Nevertheless, siderophore uptake mechanisms differ fundamentally between Gram-negative and Gram-positive bacteria, complicating community-scale investigations[21]. Gram-negative bacteria have been extensively studied: siderophores are imported via TonB-dependent outer membrane receptors (TonBDR) [22]. Structural analyses have pinpointed residues in loop L7 and the plug domain that govern receptor selectivity [23, 24], while recent bioinformatics revealed sequences near the plug domain as key determinants of pyoverdine specificity in *Pseudomonas*[25]. Building on this systematic characterization, our prior work utilized sequence coevolution between synthetase and receptor genes to reconstruct pyoverdine-mediated interaction networks in *Pseudomonas*[26]. This method assumes that the strongest coevolution signals arise between matched synthetase-receptor pairs, enabling algorithms to infer siderophore production and uptake profiles per strain. The predicted networks aligned well with experimental validations and uncovered key distinctions between pathogens and non-pathogens.

Gram-positive species lack an outer membrane and instead employ ATP-binding cassette (ABC) transporter systems to import ferric-siderophore complexes[27]. This process begins with substrate-binding proteins (SBPs), often lipoproteins tethered to the membrane, that recognize and bind siderophores with high affinity[28]. Subsequently, SBPs deliver the ligand to the permease component of the ABC transporter for ATP-driven translocation into the cytoplasm (Fig. 1a)[29]. Previous research has elucidated recognition mechanisms in model systems, including *Bacillus cereus* YxeB, which targets ferrioxamine-type siderophores via a shuttle mechanism for iron exchange[30]. However, SBPs constitute a vast and heterogeneous superfamily, classified into over 33 families in Pfam and eight structural groups, reflecting adaptations for diverse substrates including siderophores[31, 32]. Crucially, while over 1,000 unique siderophores have been characterized[8], the number of experimentally validated siderophore recognizers in Gram-positive bacteria remains disproportionately low. This vast asymmetry has prevented the definition of universal recognition motifs, leaving the molecular determinants that allow recognizers to discriminate between chemically diverse siderophores unknown. Consequently, we currently lack a predictive framework to assess the uptake potential of a Gram-positive bacterium solely from its genome, leaving a “blind spot” in the reconstruction of microbial siderophore interaction networks.

**Figure 1.**
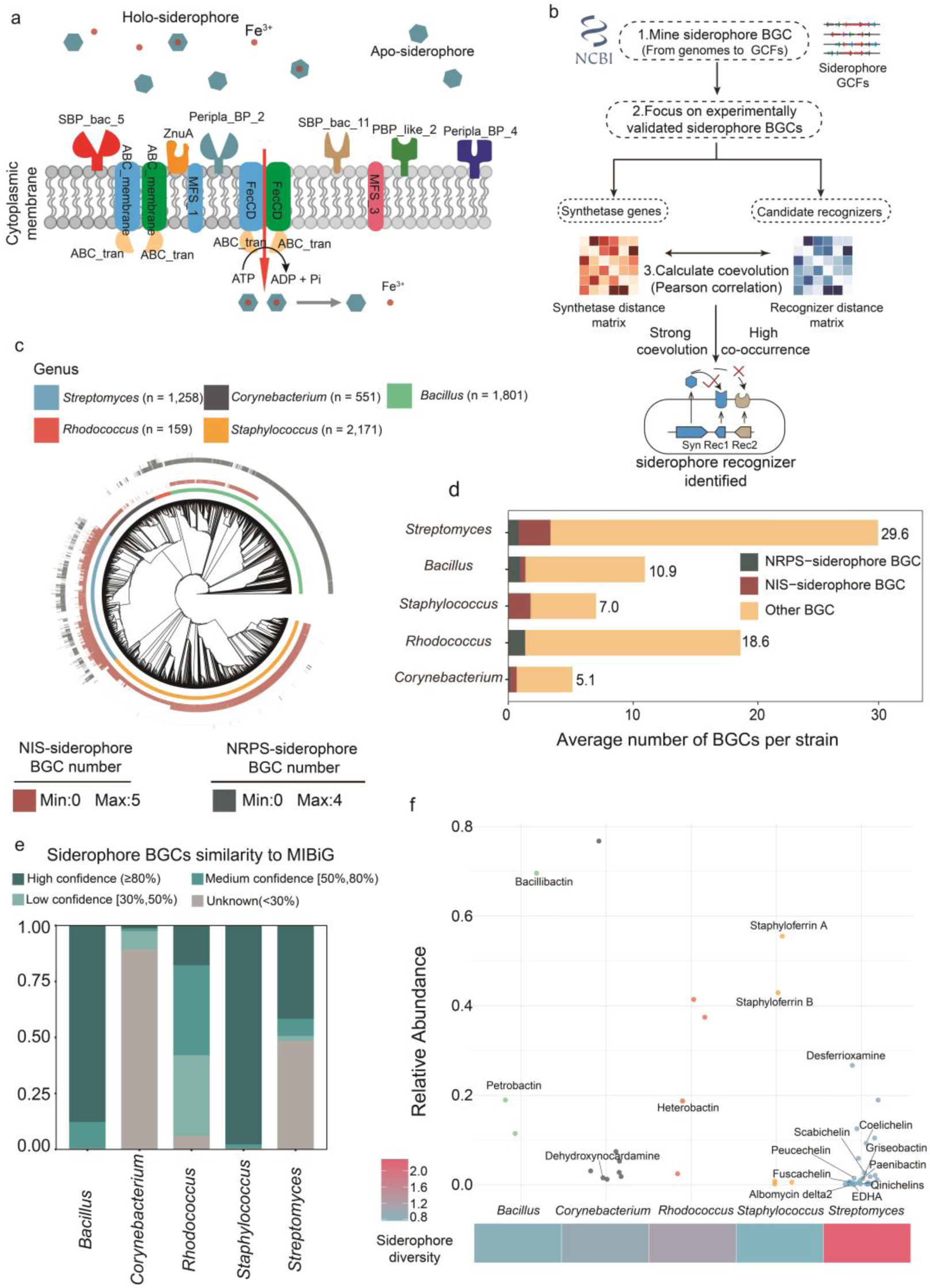
Genomic diversity and distribution of siderophore BGCs in five representative Gram-positive genera. **a.** Cartoon illustration of siderophore-mediated iron acquisition in Gram-positive bacteria. Iron-siderophore complexes are first recognized and bound by substrate-binding proteins (SBPs) on the cytoplasmic membrane, then translocated into the cytoplasm via the ABC transporter complex. Different colors and shapes denote SBP subtypes. **b.** Workflow of genome mining for the identification of potential siderophore recognizers. **c.** Phylogenetic relationships among the 5,940 strains, based on concatenated alignment of 400 single-copy conserved genes. Colors in the first ring distinguish the five genera; bar heights in the second and third rings indicate the number of NIS-type and NRPS-type siderophore BGCs per strain, respectively. **d.** BGC composition. Stacked bars show the average number of BGCs per strain across genera, color-coded as NRPS-type siderophore BGCs (green), NIS-type siderophore BGCs (red), and other BGCs (yellow). **e.** Similarity of siderophore BGCs to known BGCs in the MIBiG database. Color-coded bars highlight BGC similarity percentages against the MIBiG database: <30% (gray), [30%,50%) (light green), [50%,80%) (green), ≥80% (dark green). **f.** Relative abundance of siderophore BGCs across genera. Labeled dots denote BGCs with ≥80% similarity to MIBiG entries. Siderophore diversity was calculated using Shannon entropy, analogous to alpha-diversity in microbial communities.

In this work, we developed an integrated computational framework to bridge this knowledge gap. We systematically analyzed siderophore biosynthetic gene clusters (BGCs) and their associated uptake systems, initially focusing on five representative genera to establish ground truth, then subsequently extending our investigation to a comprehensive dataset of 16,232 Gram-positive genomes to validate universality. By integrating phylogenetic profiling, coevolutionary signal detection, and structural mapping, we sought to identify the primary siderophore recognizer and define the specific “feature sites” that drive ligand selectivity. Furthermore, we examined the genomic organization and transcriptional regulatory mechanisms governing these systems to understand how production and uptake are synchronized. This work establishes PBP2 genes as the key siderophore recognizer in Gram-positive bacteria and maps its global distribution, serving as a cornerstone for future sequence-based efforts to decode specific receptor–siderophore pairs.

## Results

### Genomic diversity and distribution of siderophore BGCs in five representative Gram-positive genera

To begin our systematic investigation of siderophore receptors in Gram-positive bacteria, we surveyed siderophore BGCs across five representative genera. These included *Bacillus* (encompassing the famous model organism *B. subtilis*[33] and pathogens *B. cereus* and *B. anthracis*[34]), *Staphylococcus* (notably *S. aureus*, a foodborne pathogen with extensively characterized iron uptake mechanisms[35]), and *Streptomyces* (known for its prolific secondary metabolism [36]). We also included *Rhodococcus* and *Corynebacterium* due to their respective industrial relevance in bioremediation and amino acid production. We collected 5,940 complete genomes that belong to these five Gram-positive bacterial genera from the NCBI RefSeq database (accessed January 21, 2025).

The phylogenetic analysis was consistent with known taxonomic classifications (Fig. 1c): Members of the class Actinomycetes (*Streptomyces*, *Rhodococcus*, and *Corynebacterium*) exhibited close phylogenetic relationships, as did *Bacillus* and *Staphylococcus* belonging to class Bacilli. AntiSMASH v7.0[37] was used to extract 78,030 BGCs. Of these, 11,095 (14.2%) were annotated as siderophore BGCs. Notably, 95.9% of the genomes contained at least one siderophore BGC (Fig. 1c). Specifically, *Staphylococcus* strains were heavily enriched in NIS-type BGCs, whereas *Rhodococcus* genomes were characterized by a predominant presence of NRPS-type clusters (Fig. 1c).

We next assessed the biosynthetic potential of the five genera (Fig. 1d). *Streptomyces* showed the highest average BGC count (29.6 BGCs per genome on average), consistent with previous reports[38], whereas *Corynebacterium* displayed the lowest (5.1 BGCs per genome on average).

We evaluated the novelty of the predicted BGCs by comparison against the manually curated Minimum Information about a Biosynthetic Gene cluster (MIBiG) database[39] (Fig. 1e). Notably, the MIBiG database exhibits genus-level biases: Well-studied genera such as *Bacillus* and *Staphylococcus* showed high similarity to the MIBiG database, with over 80% of their siderophore BGCs exhibiting similarity scores above 80% to known clusters. In contrast, less extensively studied genera like *Rhodococcus* and *Corynebacterium* had limited representation, with less than 20% of their siderophore BGCs achieving the same similarity threshold. Surprisingly, despite being among the most extensively studied genera, *Streptomyces* exhibited unexpected novelty in its siderophore biosynthetic potential, with more than half of its siderophore BGCs showing less than 50% similarity to known clusters in MIBiG. The results were cross-validated with BiG-SCAPE [40], yielded a 91.6% concordance rate (Figure S3).

To comprehensively characterize the chemical diversity of the 11,095 siderophore BGCs, we clustered all siderophore BGCs based on the pairwise sequence distance of their core biosynthetic genes (see Methods). We identified 16 types with >= 80% similarity to experimentally validated siderophores in the MIBiG database (Table S1). For the remaining NRPS-derived clusters lacking direct MIBiG hits, we annotated their chemical potential by combining antiSMASH substrate predictions with manual verification. For instance, we identified a specific class of NRPS-type siderophore BGCs in *Streptomyces* characterized by the substrate signature “diOH-Bz + Ser”, a common motif in siderophore biosynthesis (Fig. S1). Based on these siderophore BGC characterizations, we performed α-Diversity analysis in each genus. *Streptomyces* exhibited the highest diversity of siderophore BGCs (Fig. 1f).

### Identification of PBP2-subtype SBPs as the primary siderophore recognizers in representative Gram-positive genera

After characterizing the BGCs, we next sought to identify the “recognizer”, the specific membrane protein that mediates selective uptake of siderophores. Employing LLM-assisted literature mining combined with manual curation, we assembled a dataset comprising 22 experimentally validated siderophore-binding proteins localized in the cytoplasmic membrane from 11 different bacterial genera (Fig. S2a). Protein domain analysis showed that Peripla_BP_2 emerged as the only domain shared across all siderophore-binding proteins (Fig. S2b), a sensor domain in bacterial periplasmic binding proteins[41].

We extended the analysis to the five representative genera analyzed in the previous section. Previous reports showed that the Pfam database contains at least 33 substrate-binding protein (SBP) domains[31, 32]. In this study, genes encoding these domains were collectively defined as “SBP genes”. Among 11,095 siderophore BGCs from five genera, 18 of the 33 SBP domains were detected. Peripla_BP_2, Peripla_BP_3, SBP_bac_1, SBP_bac_5 and the OpuAC were among the most frequently appearing domains (Table 1). Further analysis showed that each SBP protein harbors only one SBP-related domain. Accordingly, SBPs sharing the same domain were classified into the same subtype. For clarity, we established simple abbreviations for these gene groups based on their SBP domains. For instance, full-length coding sequences containing the Peripla_BP_2 domain were designated as “PBP2 genes”. This nomenclature connects the specific conserved domain (Peripla_BP_2 domain) from the complete gene (PBP2 gene).

**Table 1:**
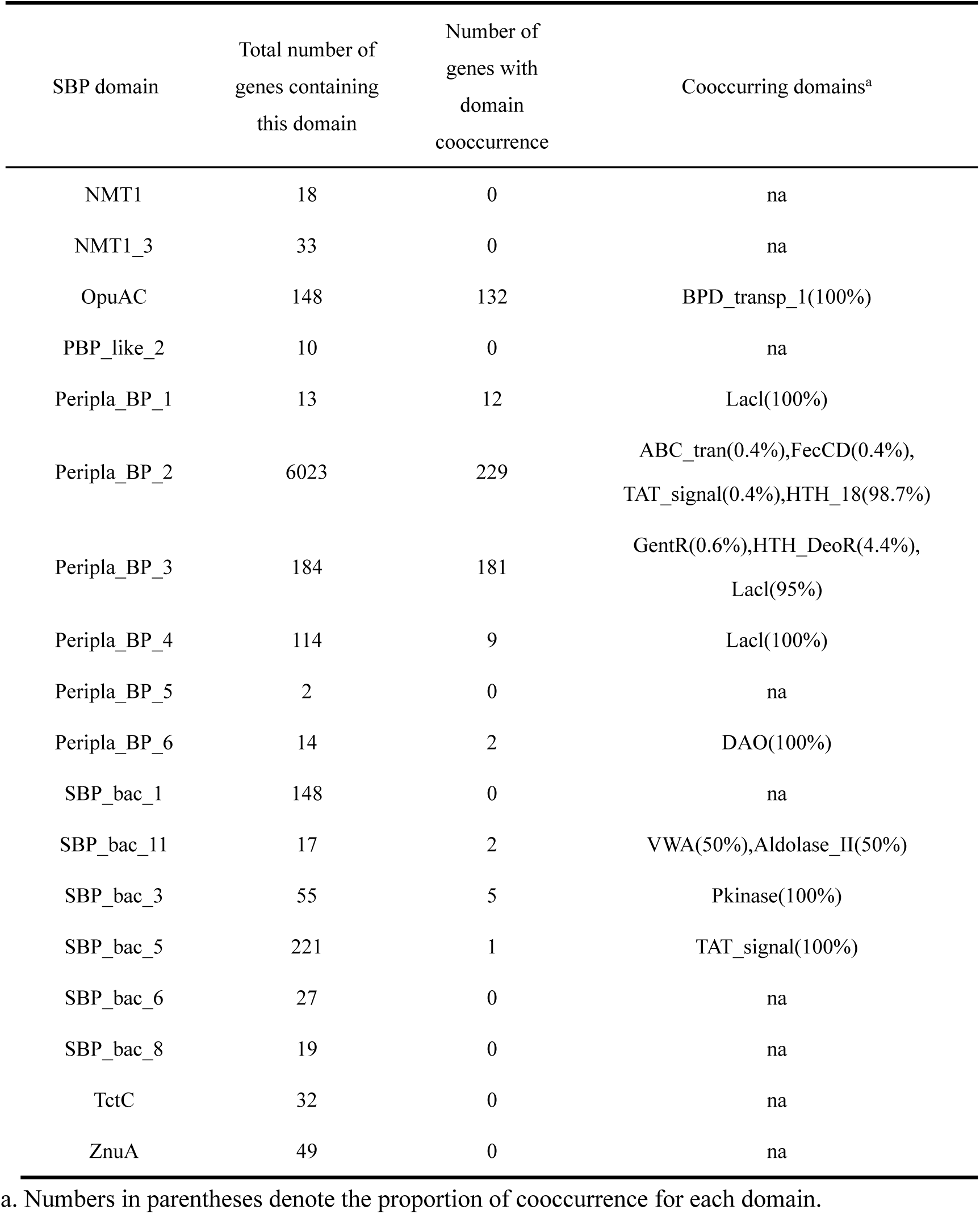
Distribution of SBP domain in siderophore BGCs and their cooccurrence with other domains.

Colocalization within a BGC was first utilized to assess coevolution (Fig. 2a, left panel), inspired by the colocalization of cognate receptors in Gram-negative siderophore BGCs[16, 42]. To ensure a balanced structural analysis of siderophore recognizers, we curated a representative dataset from the 16 experimentally characterized siderophore families identified in Section 1. We employed a stratified selection strategy to prevent analysis bias toward ubiquitous gene clusters (e.g., bacillibactin) while ensuring adequate representation of less frequent types. For each siderophore family, we selected a maximum of ten representative BGCs, prioritizing those with unambiguous biosynthetic classifications (specifically, strict “NI-siderophore” or “NRP-metallophore” types) to ensure functional integrity and exclude potential boundary artifacts. Candidates within these groups were then prioritized based on their similarity scores to the reference MIBiG cluster (see Methods). This rigorous filtering process yielded a final experimentally characterized dataset of 128 siderophore BGCs for downstream analysis.

**Figure 2.**
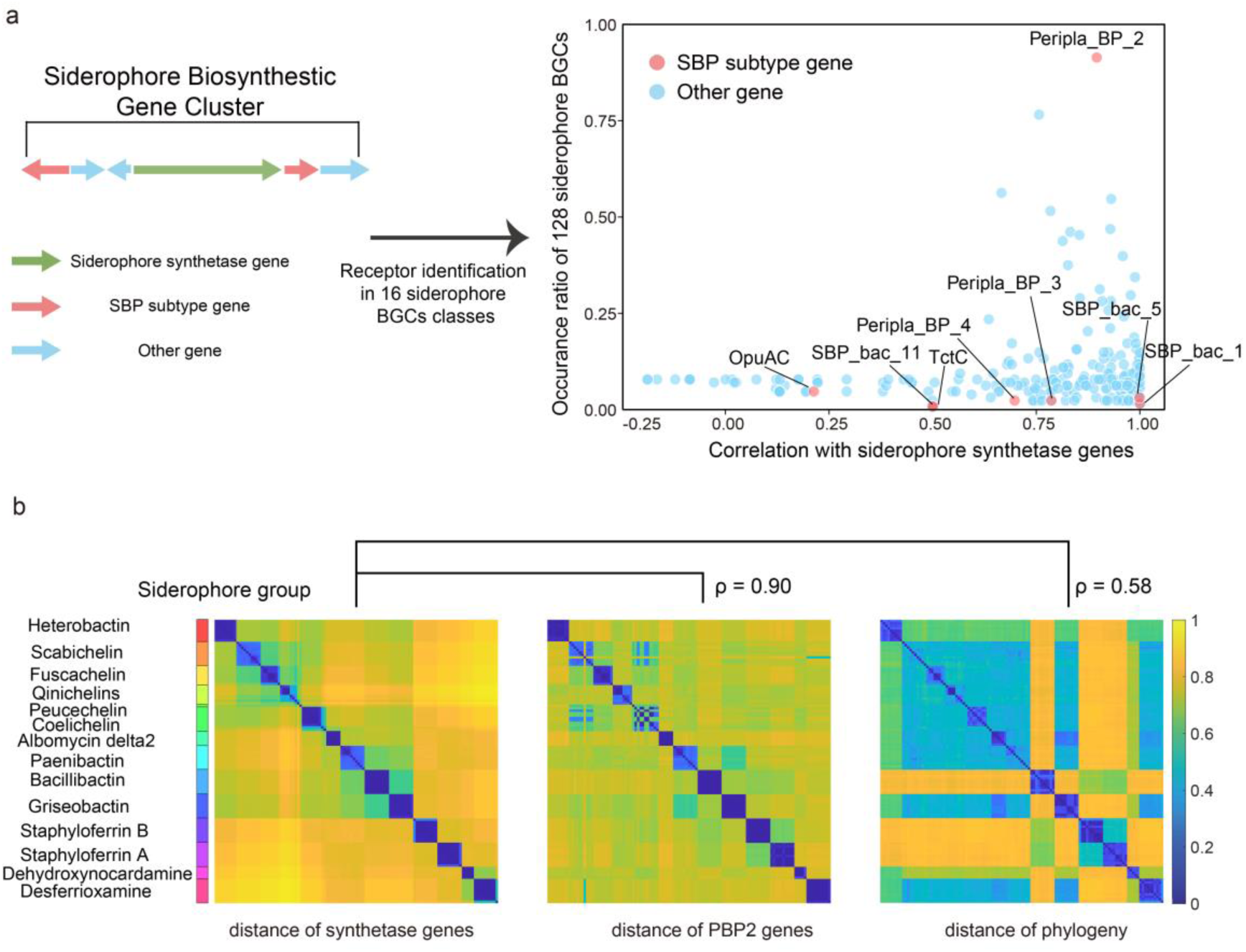
Identification of PBP2-subtype SBPs as the primary siderophore recognizers in representative Gram-positive genera. **a.** Identification of potential recognizers within 16 experimentally validated siderophore BGCs. The left panel illustrates the typical configuration of a BGC, with gene functions indicated by colors. In the right panel, each dot represents a gene class, grouped based on shared Pfam domain annotations. The x-values indicate the Pearson correlation between the sequence distance matrices of each gene class and their colocalized siderophore synthetase genes. The y-value reflects the occurrence ratio of each gene class across the siderophore BGCs. **b.** Heatmap visualizing sequence distance matrices correlations between siderophore synthetase and PBP2 genes within the same BGCs, and between siderophore synthetase genes and phylogenetic genes. This label was assigned based on high-confidence similarity (≥80%) between predicted siderophore BGCs and known clusters in the MIBiG database.

We extracted all gene classes within these 128 BGCs by their domains, excluding synthetases and calculated their occurrence frequencies (Fig. 2a, left panel, Table S2). PBP2 genes were the most prevalent class, present in 90.1% of the BGCs (Fig. 2a, right panel). To quantify coevolutionary strength[16], we calculated Pearson correlations between the sequence distance matrices of each gene class and their colocalized siderophore synthetase genes. PBP2 exhibited a strong correlation with siderophore synthetase genes (Pearson’s r = 0.90, Fig. 2a, right panel). This correlation was substantially stronger than that of the phylogenetic association (Pearson’s r = 0.58) (Fig. 2b).

### Structural characterization of specificity-determining feature sites in *Bacillus* siderophore recognizers

After identifying PBP2 as a potential siderophore recognizer, we investigated the sequence regions contributing to its specificity. To eliminate phylogenetic noise due to the substantial sequence divergence between genera, we focused on one specific genus, *Bacillus*, selected for its comprehensive set of experimentally validated receptors[19, 43]. These included FatB, FpuA, and YclQ (all three recognizing Petrobactin); FeuA (for Bacillibactin); YfiY (for schizokinen); and YxeB (for ferrichrome and desferrioxamine). Notably, while FatB and FpuA both bind Petrobactin, they target distinct moieties: FatB recognizes the 3,4-catecholate groups, while FpuA interacts with the citrate residues[39]. In contrast, YclQ predominates in non-Petrobactin-producing strains, likely serving a role in siderophore piracy[43]. Given the sequence divergence among YclQ, FatB, and FpuA, we considered these three Petrobactin recognizers as distinct PBP2 groups in our analysis.

To identify sequence regions most informative for recognizer specificity, we computed mutual information (MI) between each alignment site and the recognizer group label, with higher MI values denoting stronger associations with recognizer type. Mapping these residues to the FeuA crystal structure (PDB: 2WHY), we observed that high-MI residues were distributed across the sequence, indicating no single contiguous region dominates functional importance (Fig. 3a).

**Figure 3.**
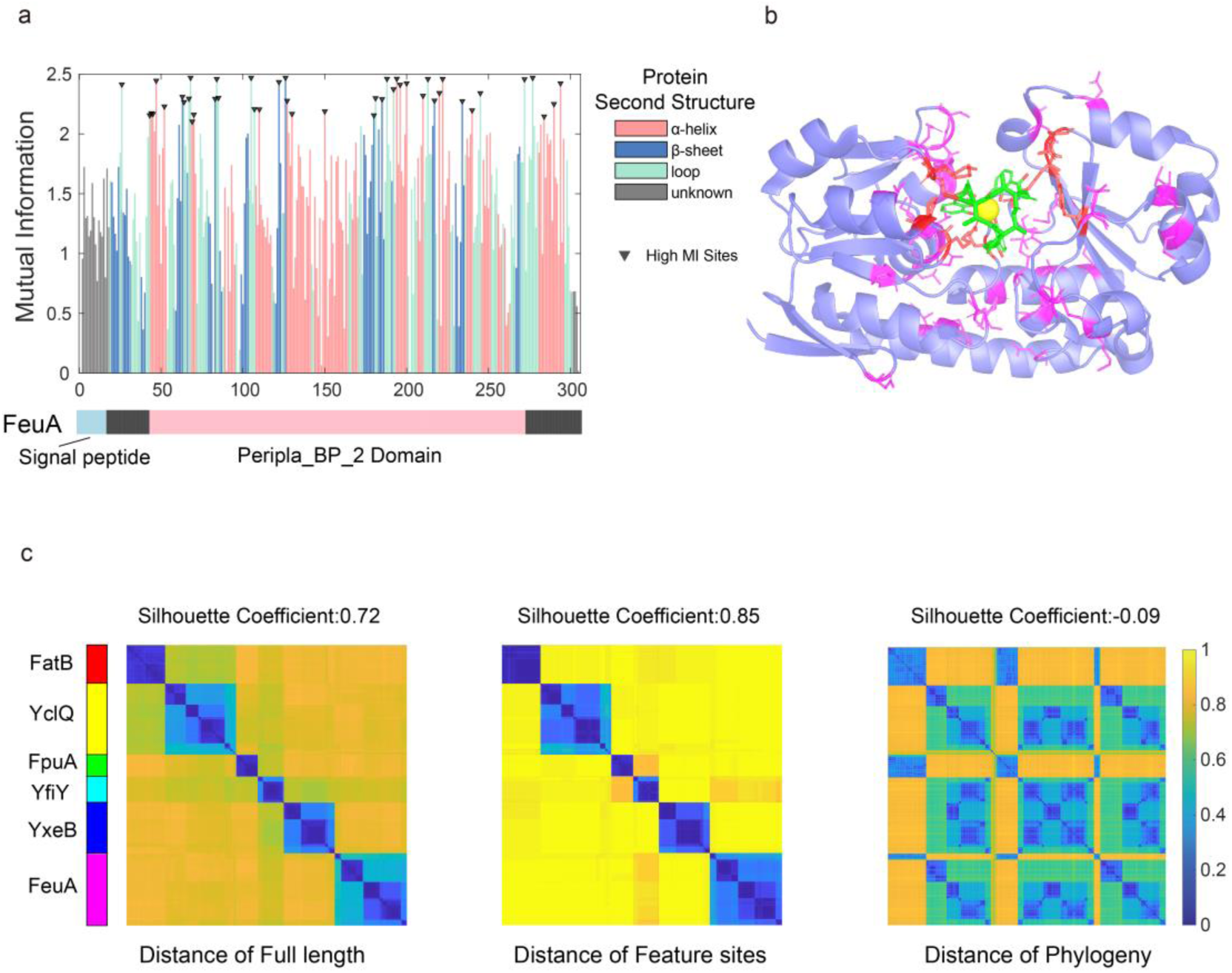
Structural characterization of specificity-determining feature sites in *Bacillus* siderophore recognizers. **a.** Mutual information analysis of 889 siderophore recognizer sequences from *Bacillus*. Residues exhibiting mutual information values exceeding 85% of the theoretical maximum MI (reflecting strong correlation with receptor grouping) are marked with black inverted triangles and defined as “feature sites.” These residues are mapped to the crystal structure of FeuA from *B. subtilis* bound to Bacillibactin (PDB: 2WHY). **b.** Crystal structure of FeuA in complex with Bacillibactin. Iron atoms are shown in yellow, Bacillibactin in green. Feature sites are highlighted in pink and red, with red indicating residues that directly interact with the ligand. **c.** Heatmaps depicting the hierarchical clustering of 889 siderophore recognizers based on pairwise distances calculated using full-length sequences (left) and feature sites only (middle). The right heatmap displays the phylogenetic distance between the corresponding strains based on concatenated alignment of 400 single-copy conserved genes. Strain clustering in all three panels is based on full-length receptor sequences. Silhouette coefficient calculated based on receptors’ label.

We benchmarked our MI scores against five residues previously confirmed by mutagenesis to be critical for FeuA–bacillibactin binding[18]. Four of the five mutagenesis-confirmed residues critical for FeuA–Bacillibactin binding (K84, K105, R180, and K213) exhibited MI values exceeding 85% of the theoretical maximum. whereas the fifth residue, R178 displayed a lower MI value. Based on the score distribution of these validated sites, we established a threshold of 85% of the theoretical maximum to define the recognizer’s “feature sites” for *Bacillus*, resulting in the identification of 44 such positions through our bioinformatic analysis.

To validate the structural role of these computationally predicted sites, we mapped them onto the experimentally solved crystal structure of the FeuA-bacillibactin complex (PDB: 2WHY). Eight of the predicted feature sites were positioned within 5 Å of the ligand, engaging in direct physical interaction (Fig. 3b).

We then compared clustering performance using full-length versus feature-site-only sequences. Pairwise sequence distances were computed, and silhouette scores were used to assess clustering quality against functional labels. Full-length sequences yielded a moderate score of 0.72. In contrast, restricting the analysis to the 44 feature sites significantly improved clustering quality to 0.85, whereas phylogeny-based clustering produced near-random partitions.

To examine the generalizability of this mechanism beyond *Bacillus*, we extended our structural analysis to *Streptomyces*, the genus with the highest siderophore biosynthetic potential in our dataset. By curating a high-confidence dataset of 980 *Streptomyces* PBP2 sequences covering 12 distinct siderophore types and using the DesE crystal structure (PDB: 6ENK) as a reference, our Mutual Information analysis identified 21 feature sites that substantially improved the resolution of distinct siderophore specificities compared to full-length sequences (Figure S4). Consistent with our *Bacillus* results, these sites spatially cluster around the ligand-binding pocket (Figure S5).

### Genomic architecture and transcriptional regulation of PBP2-mediated iron uptake in representative genera

Confident that PBP2 genes serve as primary siderophore recognizers, we examined their genome-wide distribution across the 5,940 genomes. HMMER screening revealed PBP2 genes exist in all genomes, ranging from 1 to 49 copies per genome, with over 90% of strains harboring 4-12 copies (Fig. 4a). Notably, *Rhodococcus erythropolis* and *Rhodococcus qingshengii* stand out with an exceptionally high average of 46.37 PBP2 genes per genome (Fig. 4a). These two species are commonly found in iron-limited environments such as arsenic-contaminated soil, Antarctic soil, and weathered serpentine rocks [44–46].

**Figure 4.**
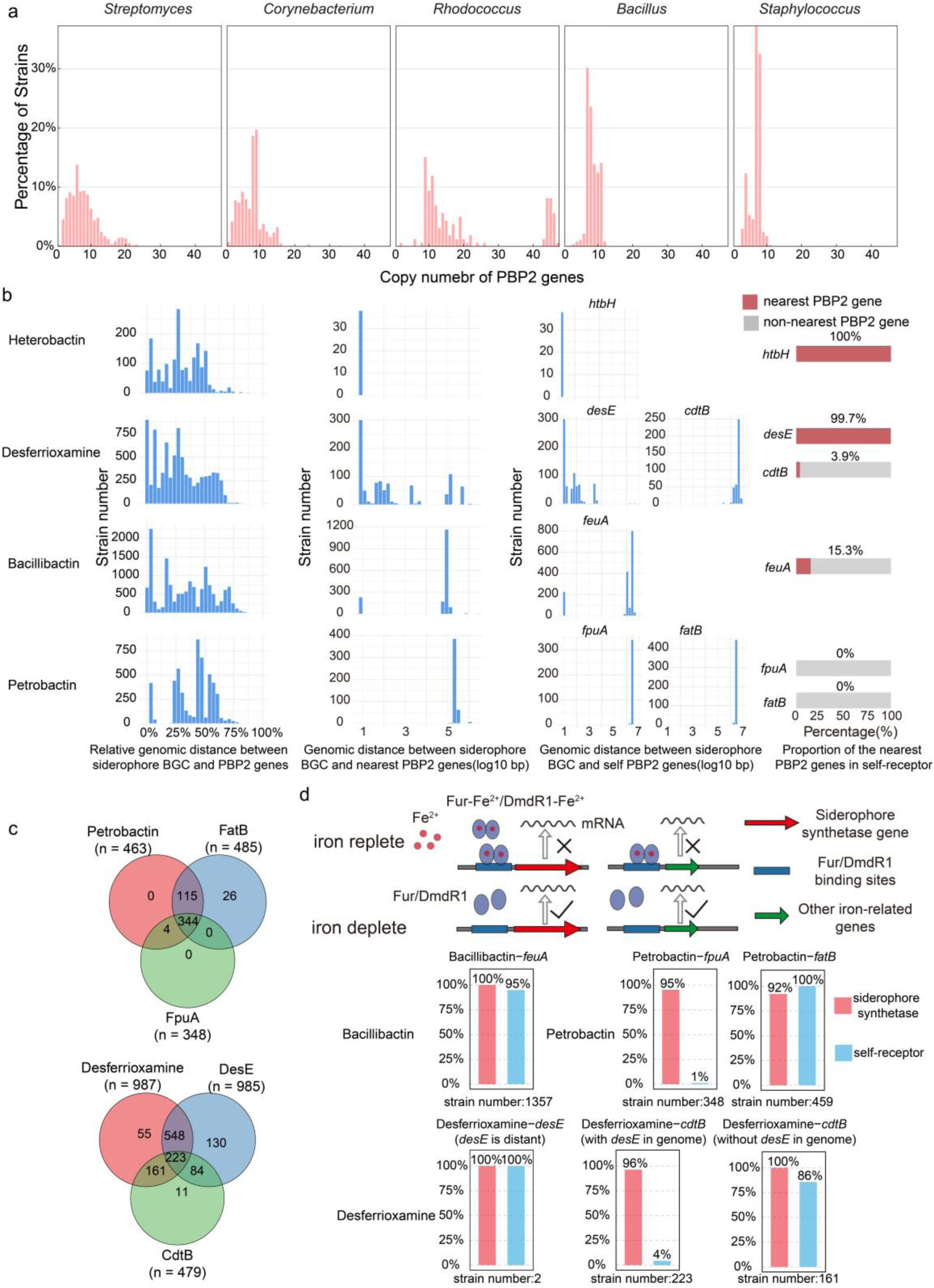
Genomic architecture and transcriptional regulation of PBP2-mediated iron uptake in representative genera. **a.** Distribution of PBP2 gene number among five genera. The x-axis represents the number of PBP2 genes, while the y-axis shows the number of strains. **b.** From left to right: (i) genomic distribution of all PBP2 genes with siderophore BGCs set as the reference points; (ii) distribution of the genomic distances between siderophore BGCs and their nearest PBP2 genes (log10 scale); (iii) distribution of the genomic distances between self-receptors and siderophore BGCs (log10 scale); and (iv) proportion of self-receptors that are also the nearest PBP2 genes to siderophore BGCs. **c.** Venn diagram showing the overlap between strains harboring siderophore biosynthetic genes and those harboring receptor genes. **d.** Proportion of siderophore synthetase and receptor genes regulated by Fur/DmdR1 protein. The diagram illustrates the Fur/DmdR1 protein regulation mechanism.

Inspired by receptor-BGC colocalization in Gram-negative bacteria, we analyzed PBP2 genomic positions relative to siderophore BGCs, uncovering three patterns(Fig. 4b): (1) Consistent colocalization, where every BGC of a type includes a PBP2 receptor (e.g., Heterobactin in *Rhodococcus*); (2) Partial colocalization, where only some BGCs contain a PBP2 gene (e.g., ∼78% of Desferrioxamine BGCs, and ∼15% of Bacillibactin BGCs); and (3) Complete absence, where the BGC completely lacks a PBP2 gene within the cluster (e.g., Petrobactin BGCs).

Desferrioxamine, a crucial siderophore in *Streptomyces*, exists as linear Desferrioxamine B and cyclic Desferrioxamine E. Both DesE and CdtB receptors recognize these forms, though DesE exhibits ∼100-fold higher affinity for Desferrioxamine E than CdtB[47]. Among the 5,940 analyzed genomes, 987 harbor a Desferrioxamine BGC, with 771 also containing *desE* gene. The *desE* gene typically colocalizes with the Desferrioxamine BGC (within 6 kb), with only two genomes showing a distance >1 Mb. In contrast, *cdtB* is encoded in only 384 Desferrioxamine - producing genomes and never clearly colocalizes with the Desferrioxamine BGC (Fig. 4b-c). For Petrobactin BGC, *fatB* and *fpuA* are generally >1 Mb from the BGC (Fig. 4b). Nearly all Petrobactin-containing strains encode *fatB*, whereas only ∼75% carry *fpuA* (Fig. 4c). These contrasting genomic distributions of siderophore BGCs and their cognate recognizers prompt an investigation into their coregulation.

Ferric uptake regulator (Fur in *Bacillus*) and divalent metal-dependent regulatory protein 1 (DmdR1 in *Streptomyces*) have been experimentally confirmed to regulate iron homeostasis, yet the genomic distributions of their target genes have not been investigated thoroughly[48–50]. Using position weight matrix (PWM) based scanning, we identified potential Fur/DmdR1 binding sites within BGCs and upstream of recognizer genes. For Bacillibactin, 1,356/1,357 BGCs and 1,288 *feuA* loci harbor Fur sites, indicating coregulation. For Petrobactin (459 strains), 92% of BGCs and all *fatB* promoters possessed Fur sites, but only 1% of *fpuA* loci did. For Desferrioxamine, *desE* is typically within the BGC, facilitating coregulation. In the only two distant (>1 Mb) *desE* cases, both *desE* and the Desferrioxamine BGC remain DmdR1-regulated. Conversely, *cdtB* showed weaker DmdR1 association: ∼4% coregulation when *desE* is present. Strikingly, in *desE*’s absence strains, 86% of *cdtB* loci and all Desferrioxamine BGCs are DmdR1-controlled.

### Universality of PBP2 genes and diverse iron acquisition strategies across Gram-positive bacteria

To broaden the taxonomic scope of our initial analysis, we expanded the investigation to encompass a comprehensive dataset of Gram-positive bacteria. The dataset spans the phyla Bacillota, Actinomycetota, and Chloroflexota, thereby covering the majority of Gram-positive lineages[51]. By filtering for monoderm lineages and excluding strains encoding the outer membrane assembly protein BamA[52], we curated a dataset of 16,232 genomes (see Methods).

PBP2 genes exhibited a pervasive distribution within this expanded collection. Specifically, 14,197 (87.5%) of the strains harbored at least one PBP2 gene (Fig. 5a). Intriguingly, analysis of the 2,035 genomes lacking PBP2 genes revealed that they predominantly cluster into three specific ecological or physiological categories that do not constrain tightly by iron (Fig. S6): (1) lactic acid bacteria (e.g., *Lactobacillus*, *Leuconostoc*) that utilize manganese as an iron substitute; (2) obligate anaerobes (e.g., *Clostridium*, *Bifidobacterium*) residing in iron-replete niches rich in ferrous iron; and (3) Mollicutes (e.g., *Mycoplasma*, *Spiroplasma*) possessing highly reduced genomes dependent on host-derived nutrients.

**Figure 5.**
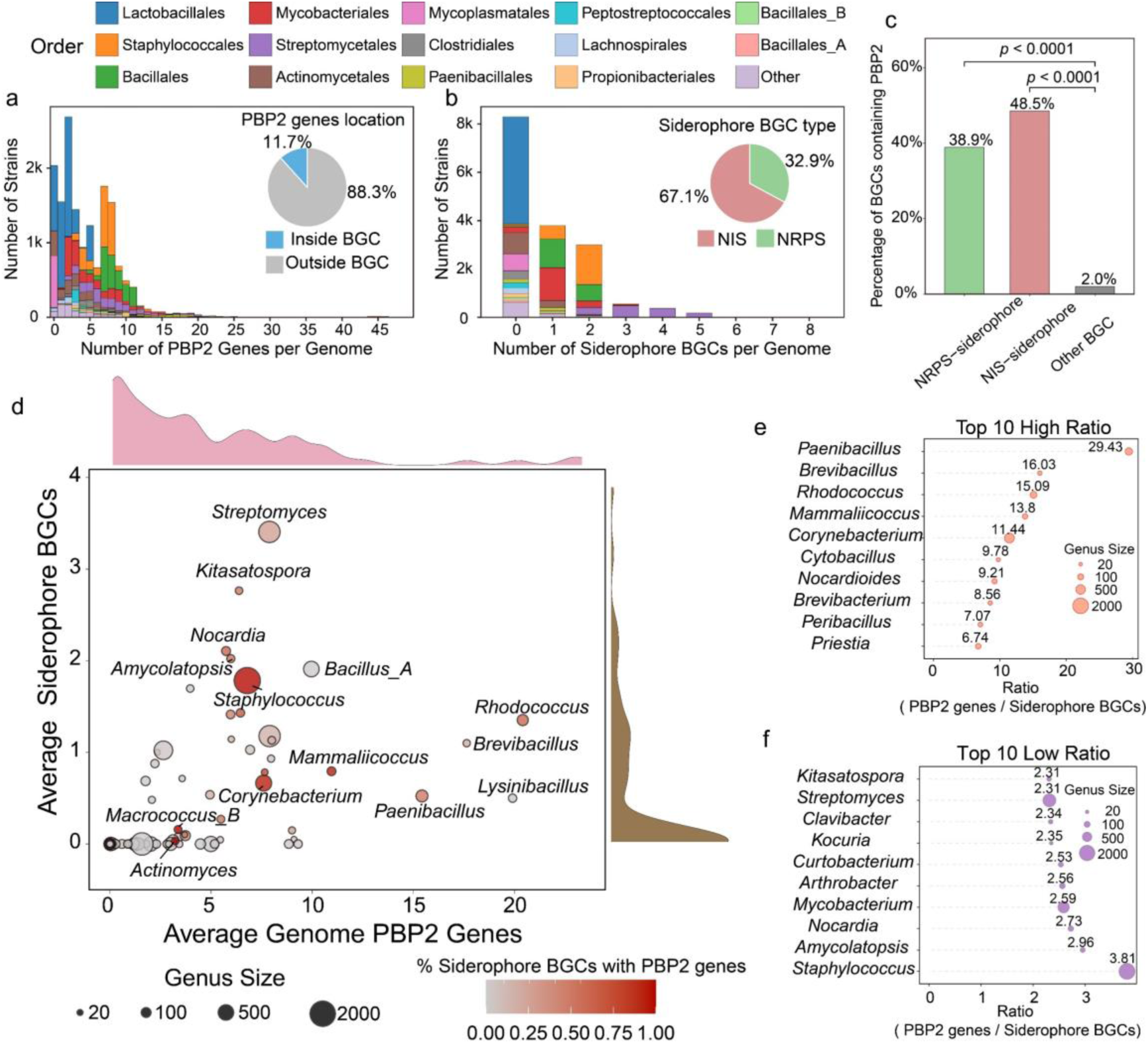
Universality of PBP2 as a siderophore recognizer and divergence of iron acquisition strategies across Gram-positive bacteria. **a.** Distribution of PBP2 gene counts per genome across 16,232 genomes, stacked by taxonomic order (colors correspond to the legend at the top). The inset pie chart displays the genomic location of PBP2 genes (Inside versus Outside BGCs). **b.** Distribution of siderophore BGC counts per genome across 16,232 genomes, stacked by taxonomic order. The inset pie chart illustrates the proportion of siderophore biosynthetic types (NRPS versus NIS). **c.** Occurrence of PBP2 genes within siderophore BGCs compared to other BGCs. The bar chart shows the percentage of BGCs containing PBP2 genes across NRPS-siderophore, NIS-siderophore, and Other BGC categories, indicating a significant functional coupling between PBP2 genes and siderophore BGCs (p < 0.0001, pairwise Fisher’s exact test). **d.** Genus-level patterns of siderophore synthesis and uptake potential. Each dot represents a genus, plotted by the average number of PBP2 genes (uptake potential, x-axis) versus the average number of siderophore BGCs (synthesis potential, y-axis) per genome. Point size indicates the number of genomes analyzed in each genus, and the color gradient represents the proportion of siderophore BGCs that contain internal PBP2 genes. Marginal density plots show the distribution of each variable. **e-f.** Genera with the highest (e) and lowest (f) ratios of total genomic PBP2 genes to siderophore BGCs. Dots are sized by the number of genomes in the genera. This analysis was restricted to genera with an average of >0.5 siderophore BGCs per genome to highlight significant evolutionary strategies.

We next evaluated the prevalence of siderophore BGCs. A total of 7,933 (48.9%) genomes contained predicted siderophore BGCs (Fig. 5b), with a distinct enrichment in the order Actinomycetales. Within these identified clusters, the NIS pathway accounting for 67.1% of the total siderophore BGCs, compared to 32.9% for the NRPS type (Fig. 5b).

To assess the functional coupling within these systems, we quantified the cooccurrence of PBP2 genes and siderophore BGCs in each genome. The total count of identified PBP2 genes (79,337 counts) significantly outweighed that of siderophore BGCs (13,999 counts), with the vast majority of PBP2 genes (88.3%) located outside of siderophore BGCs (Fig. 5a). However, from the perspective of BGC composition, we found that 38.9% of NRPS-siderophore BGCs and 48.5% of NIS-siderophore BGCs contained at least one PBP2 gene (Fig. 5c). These frequencies were significantly higher than the background rate observed in other functional BGC categories (2.0%; p < 0.0001, pairwise Fisher’s exact test) (Fig. 5c). Furthermore, a comprehensive analysis of SBP genes within all predicted siderophore BGCs confirmed that PBP2 genes constituted the most abundant SBP subtype genes (Fig. S7).

We subsequently quantified the patterns of siderophore synthesis and uptake at the genus level (Fig. 5d). Regarding biosynthetic potential, genera such as *Streptomyces* and *Kitasatospora* have the highest numbers of siderophore BGCs (Fig. 5d, Fig. S8). In terms of uptake potential *Rhodococcus* and *Lysinibacillus* possessed the most PBP2 genes (Fig. 5d, Fig. S8). If we quantify genomic coupling between siderophore BGCs and PBP2 genes by the proportion of BGCs containing a PBP2 gene, we found that the degree of coupling varied considerably even among phylogenetically related genera (Fig. 5d). For example, *Corynebacterium* displayed a high degree of coupling (93%), contrasting with substantially lower proportions in the related genera *Streptomyces* (59%) and *Rhodococcus* (26%). To quantify the balance between siderophore synthesis and uptake, we calculated the ratio of genomic PBP2 genes to siderophore BGCs for genera with an average of >0.5 siderophore BGCs per genome (Fig. 5d). *Paenibacillus* displayed the highest ratio (29.43). Conversely, *Nocardia* and *Mycobacterium* showed some of the lowest ratios (<3.0) (Fig.5e, f).

## Discussion

The ability to infer microbial interactions from genomic data has revolutionized our understanding of community dynamics, particularly for the well-characterized iron acquisition systems of Gram-negative bacteria[4]. However, a parallel understanding for Gram-positive bacteria has remained elusive due to the lack of a defined siderophore receptor, creating a significant blind spot in predictive microbial ecology. In this study, we first mined 5,940 genomes from five key Gram-positive genera to pinpoint PBP2subtype SBP as the main siderophore recognizer. We demonstrate that PBP2 exhibits a strong coevolutionary signal with synthetase genes that transcends phylogenetic relatedness, establishing PBP2 not merely as a frequent genomic neighbor (90.1% cooccurrence ratio) but as the coevolving functional partner required for uptake.

Expanding our scope to 16,232 genomes covering most Gram-positive bacteria confirmed PBP2 as a universal recognizer, present in 87.5% of monoderm bacteria. Intriguingly, the exceptions reinforce the rule. The 12.5% of genomes lacking PBP2 genes predominantly clustered into lineages that do not rely on ferric-siderophore uptake, such as: manganese-utilizing Lactobacillales or obligate anaerobes residing in ferrous-rich environments The absence of PBP2 in these lineages—where iron selective pressure is relaxed—reinforces the conclusion that PBP2 is strictly maintained only when siderophore-mediated iron acquisition is ecologically essential. Furthermore, the quantitative balance between uptake and biosynthetic genes highlights distinct life-history strategies across genera. We observed that genera such as *Paenibacillus* possess a remarkably high receptors-to-synthetase ratio (∼29:1), indicative of a “scavenging” lifestyle focused on pirating xenosiderophores. Similarly, *Rhodococcus* species have a large amount of PBP2 genes (up to 46 per genome), likely an adaptation to capture diverse iron sources in oligotrophic environments. In contrast, pathogens like *Mycobacterium* maintain a lower receptors-to-siderophore BGCs ratio (<3:1), reflecting a “self-sufficient” strategy. Consequently, PBP2 gene counts offer a genomic signature to predict whether a bacterium functions primarily as a cooperator or a cheater in iron-limited communities.

The first notable difference between Gram-positive and Gram-negative siderophore recognition systems lies in the arrangement of specificity-determining sites. In Gram-negative bacteria, these sites cluster in the plug domain and loop L7 of TonBDR [23–25]. In contrast, our analysis reveals that PBP2 feature sites are scattered across the sequence. This sequence-level dispersion is intrinsic to the “Venus flytrap” mechanism of SBPs, where two globular domains are connected by a flexible hinge close around the ligand[32]. Consequently, residues that are distant in sequence are brought into spatial proximity to form the binding cleft, naturally resulting in a discontinuous interface. Evolutionarily, the absence of a tight binding hotspot in PBP2 may reflect a more flexible recognition mechanism involving allosteric effects or conformational shifts, enabling promiscuity. This interpretation aligns with previous case reports of crossover recognition in Gram-positive bacteria: in *Bacillus subtilis*, FeuA binds and imports both its cognate catecholate siderophore bacillibactin and the structurally similar xenosiderophore enterobactin[18, 53], while in *Bacillus cereus*, FpuA and FatB interact with both apo- and ferric-petrobactin[27, 30].A second divergence appears in genomic organization. In Gram-negative bacteria like *Pseudomonas*, cognate TonBDRs are typically colocalized with siderophore biosynthetic gene clusters to ensure the simultaneous inheritance of all related genetic elements via horizontal gene transfer [16, 42]. Conversely, Gram-positive systems often exhibit spatial decoupling, where PBP2 genes often lie distant from their cognate BGCs yet maintain functionally coordination through shared iron regulators like Fur or DmdR1. This spatial divergence may reflect lower HGT rates in Gram-positive lineages, which relax the pressure for physical clustering and allow for transcriptional rather than physical coupling. [54–56]. This spatial flexibility enables Gram-positives to mix receptors across clusters for enhanced versatility. For instance, in desferrioxamine pathways, the primary receptor DesE resides within the biosynthetic cluster, while the distant, promiscuous backup receptor CdtB ensures uptake even if the biosynthetic locus is lost or mutated [47]. Such distributed, redundant networks buffer Gram-positive bacteria against environmental fluctuations, contrasting with the rigid, all-or-nothing organization of Gram-negative systems.

Phylogenetic analyses suggest that Gram-positive and Gram-negative lineages diverged approximately 2.2 to 3.2 billion years ago—significantly predating the Great Oxidation Event (GOE)[57]. Consequently, when the GOE subsequently triggered a global iron crisis by oxidizing ferrous iron[58], these two lineages were already evolutionarily distinct. This timing implies that siderophore-mediated iron acquisition evolved as independent solutions to a shared selection pressure[59]. Therefore, these distinct recognizer properties in Gram-positive and Gram-negative systems reflect separate evolutionary paths and selection pressures.

Despite these insights, our study encompasses several limitations. First, while our genomic abundance survey spanned the full breadth of Gram-positive diversity (∼16,000 genomes), our fine-grained coevolutionary analysis was necessarily restricted to a curated dataset of 128 BGCs from five representative genera. This stratified downsampling was compelled by the extreme data imbalance in public databases, where model organisms heavily predominate. Second, although our bioinformatic predictions of “scattered” feature sites align with the “Venus flytrap” structural mechanism, experimental validations such as binding assays or crystallography are required to definitively confirm the functional roles of these residues and the predicted promiscuity. Finally, our inferences regarding transcriptional synchronization relied on Position Weight Matrix (PWM) scanning of known Fur/DmdR1 motifs, which may miss non-canonical binding sites or novel species-specific regulatory elements.

Looking ahead, our systematic characterization of PBP2 deepens insights into how Gram-positive and Gram-negative bacteria independently adapted their iron acquisition strategies to post-GOE iron scarcity. Moreover, by establishing PBP2 as the primary siderophore recognizer, this work bridges a critical gap in Gram-positive iron acquisition. This foundational identification enables the global mapping of uptake potential across diverse taxa, paving the way for genome-based inference of siderophore-mediated interaction networks in complex microbial communities.

## Methods

### Data collection and taxonomy detection

A total of 46,362 complete genomes were retrieved from the NCBI RefSeq database (RRID:SCR_003496) as of January 21, 2025. Taxonomic assignments were performed using the Genome Taxonomy Database (GTDB, release R220, RRID:SCR_019136) with default parameters. Among these genomes, 1,258 were classified as *Streptomyces*, 159 as *Rhodococcus*, 551 as *Corynebacterium*, and 2,171 as *Staphylococcus*. The classification of *Bacillus* was treated collectively because GTDB subdivides the traditional Bacillus genus (including *B. subtilis*, *B. cereus*, *B. anthracis*, and *B. thuringiensis*) into multiple lineages that retain the prefix “*Bacillus*”. To ensure consistency, all GTDB-defined genera beginning with “*Bacillus*” were aggregated and analyzed together, totaling 1,801 genomes.

### Construction of phylogenetic tree

A phylogenetic tree (Fig. 1c) was constructed using the PhyloPhlAn3 pipeline (RRID:SCR_013082), which integrates marker selection, multiple sequence alignment, and phylogenetic inference. Specifically, the analysis was executed using the “supermatrix_aa.cfg” configuration file with the “--diversity medium” parameter to ensure accurate resolution across the analyzed genera. Within this automated pipeline, universal single-copy orthologs (>400 marker genes) were identified using Diamond, followed by multiple sequence alignment using MAFFT. Poorly aligned regions were rigorously trimmed using trimAl. The final maximum likelihood phylogeny was inferred using RAxML. The resulting tree topology was visualized and annotated using the R package “ggtree” and “ggtreeExtra”.

### Clustering analysis to group siderophore biosynthetic genes

All core biosynthetic genes were extracted from siderophore BGCs. Pairwise sequence distances were computed as p-distances (the proportion of differing residues between two aligned sequences) using the “seqpdist” function in MATLAB (R2024a; RRID:SCR_001622) with the “PairwiseAlignment” option set to false. Hierarchical clustering was applied to group the genes, and the optimal clustering threshold was determined based on silhouette scores and the resulting number of clusters (Figure S1). To minimize phylogenetic bias, clustering was conducted independently for each genus.

### Genome mining to detect BGCs and annotated siderophore BGCs

Genome mining was performed using antiSMASH v7.0.0 (RRID: SCR_022060) on annotated genomes to identify secondary metabolite biosynthetic gene clusters. The analysis was run with the following parameters: “--asf --clusterhmmer --cc-mibig –tfbs --cb-knownclusters --cb-subclusters”. Clusters classified as either “NRP-metallophore” or “NIS-siderophore” were designated as siderophore BGCs, corresponding to NRPS-type and NIS-type pathways, respectively (Fig. 1c, d). The similarity between predicted BGCs and known entries in the Minimum Information about Biosynthetic Gene cluster (RRID:SCR_023660) was evaluated using the built-in “***knownclusterblast”*** function, with a similarity cutoff of ≥80% for tentative annotation as known metabolites (Fig. 1e).

### Siderophore BGC novelty analysis

To systematically assess the novelty of the identified siderophore BGCs, we employed a dual-metric approach combining sequence similarity (antiSMASH) and gene cluster architecture distance (BiG-SCAPE). Initial screening was performed using the antiSMASH “ ***knownclusterblast*** “ module, where BGCs were classified as “known” if they exhibited >=80% similarity to an entry in the MIBiG database. To cross-validate these findings using domain-based architectural similarity, a global network analysis was executed using BiG-SCAPE (v2.0.0) against the MIBiG database (v4.0). The analysis utilized the --mix parameter to enable clustering across different BGC classes and --alignment-mode local to optimize domain alignment, with Gene Cluster Family (GCF) cutoffs set to 0.3 including singletons. The resulting raw network output was processed using a custom Python script to extract the “best MIBiG hit” for each query, defined as the reference cluster exhibiting the minimum global alignment distance. A strict novelty threshold was established for BGCs with a minimum distance > 0.3 to their nearest MIBiG neighbor, as this cutoff demarcates distinct chemical families.

### Construction of an experimentally characterized, taxonomically balanced dataset Siderophore BGC Dataset

To facilitate unbiased structural and co-evolutionary analyses, we curated a representative dataset based on the 16 experimentally characterized siderophore families identified in our initial survey. For each siderophore family, a maximum of ten representative BGCs were selected based on a hierarchical prioritization scheme. First, to ensure functional integrity and exclude potential boundary artifacts, selection was restricted to BGCs with unambiguous biosynthetic classifications (specifically, strict ‘NI-siderophore’ or ‘NRP-metallophore’ types). Second, candidates within these categories were prioritized according to their sequence similarity scores to the reference MIBiG cluster. This curation process yielded a final dataset comprising 128 high-confidence siderophore BGCs for downstream analysis.

### Calculation of the correlation between synthetase and candidate genes located within siderophore BGC sequence distance matrices

To identify potential siderophore receptors, we quantified the coevolutionary relationships between genes located within siderophore BGCs and their corresponding biosynthetic genes. We first performed hierarchical clustering of the siderophore biosynthetic genes based on sequence distance. Using the same order derived from this clustering, we then reordered the pairwise sequence distance matrix (calculated using p-distance) for all candidate genes within the BGCs. Finally, to evaluate the coevolutionary association between the two matrices, the parametric Pearson correlation coefficient was calculated using the “corr” function in MATLAB. (R2024a; RRID:SCR_001622).

### Collection and domain analysis of known siderophore SBPs

Twenty-two experimentally validated siderophore SBPs were curated from the literature. Protein sequences were obtained from the Protein Data Bank (PDB; RRID:SCR_012820) and NCBI (RRID:SCR_006472). Multiple sequence alignment was conducted using Clustal Omega (RRID:SCR_001591) with default parameters. A phylogenetic tree was inferred using the Neighbor-Joining method based on pairwise amino acid distances calculated under the WAG substitution model and was midpoint-rooted. Conserved domains were identified using “hmmscan” from HMMER v3.0 (RRID:SCR_005305) against the Pfam database (RRID:SCR_004726), retaining all hits with a bit score > 0, and the domain architectures shared among these SBPs were analyzed.

### Identification of siderophore receptors, multiple sequence alignment, and detection of feature sites in *Bacillus*

Six experimentally characterized siderophore receptors from *Bacillus* were collected from published literature. Homologous sequences were identified across 1,801 *Bacillus* genomes using BLASTP (RRID:SCR_001010) with an “E-value” cutoff of 1e-5, yielding 889 unique receptor sequences after redundancy removal. Multiple sequence alignment was performed using Clustal Omega (RRID:SCR_001591) with default parameters. Mutual information (MI) between alignment positions and receptor identity labels was calculated to assess residue specificity. Secondary structure information was derived from the FeuA crystal structure using DSSP (RRID:SCR_016067) with default parameters.

### Construction of high-confidence siderophore recognizer dataset and feature site analysis in *Streptomyces*

Genome mining across 1,258 *Streptomyces* genomes initially identified 4,222 siderophore BGCs using antiSMASH. To establish a high-confidence dataset of siderophore recognizers, PBP2 genes were first identified using “hmmsearch” (E-value < 1e-5) and subsequently filtered based on genomic context. Only PBP2 genes physically colocalized within annotated siderophore BGCs were retained. To eliminate potential ambiguity in substrate assignment, the dataset was further restricted to BGCs containing exactly one PBP2 gene. This automated selection process captured the experimentally validated receptors DesE and CchF, which are encoded within their respective siderophore BGCs. The dataset was manually supplemented with the experimentally characterized receptor CdtB, which is typically encoded outside of siderophore BGCs. The final curated dataset comprised 980 unique PBP2 sequences covering 12 distinct siderophore classes. Feature sites were identified using Mutual Information (MI) analysis as described for *Bacillus*, utilizing the DesE crystal structure (PDB: 6ENK) as the structural reference. Given the high sequence diversity observed in this genus relative to *Bacillus*, the threshold for defining feature sites was set at 80% of the theoretical maximum MI. The resolution of functional specificity was evaluated by calculating Silhouette coefficients using pairwise distances derived from feature-site identity compared to full-length sequence identity.

### Genome-wide identification of PBP2 genes

Protein-coding sequences ranging from 200 to 500 amino acids were screened using “hmmsearch” from HMMER (RRID:SCR_005305) against the PBP2 domain profile (PF01497) from the Pfam database (RRID:SCR_004726). Proteins with significant hits (E-value < 1e-5) were retained as putative PBP2 gene candidates.

### Identification of Fur/DmdR1 binding sites upstream of genes

Nineteen experimentally validated Fur-binding sites from Bacillus subtilis were used to construct a position weight matrix (PWM) using the MEME Suite v5.5.7 (RRID:SCR_001783). In parallel, the PWM for DmdR1 was obtained from the LogoMotif database(https://logomotif.bioinformatics.nl/). The Fur PWM was used by FIMO (RRID:SCR_001783) to scan the upstream regions of all genes across *Bacillus* genomes, whereas the DmdR1 PWM was applied to all *Streptomyces* genomes. Motif occurrences with p < 0.001 were considered significant.

### Construction of a rigorous Gram-positive (monoderm) bacterial dataset

To strictly define the Gram-positive (monoderm) dataset and exclude potential diderm contaminants, we applied a two-step filtration strategy combining taxonomic and proteomic criteria. First, taxonomic filtering: Genomes belonging to the phyla *Bacillota*, *Actinomycetota*, and *Chloroflexota* were selected based on GTDB (GTDB, release R220, RRID:SCR_019136) metadata, while the class *Negativicutes* (a known diderm lineage within *Bacillota*) was explicitly excluded. Second, proteomic screening for outer membrane markers: To eliminate cryptic diderm lineages that might be misclassified, we screened all proteomes for the presence of BamA (Omp85), an essential outer membrane assembly factor (Pfam: PF01103). The identification of BamA were conducted by “hmmsearch” with a stringent E-value cutoff of 1e-5. To prevent false positives derived from protein fragments, only hits with a sequence length between 300 and 1,500 amino acids were considered valid. Strains possessing significant BamA hits were classified as diderms and were excluded from the final dataset, ensuring that only true monoderm bacteria were retained for downstream analysis.

### Data, code, and reproducibility

All genomes analyzed in this study are publicly available through the NCBI RefSeq database (RRID:SCR_003496). The full list of genome accessions is provided in Supplementary Dataset 1. All computational analyses were performed using version-controlled scripts and software with fixed random seeds to ensure reproducibility. The complete workflow, including data processing and analysis steps illustrated in Figure 1b, together with all associated scripts, is available on GitHub: https://github.com/Linlong-Yu/PBP2-as-the-Predominant-Siderophore-Recognizer and the Figshare portal. The full computational protocol is also accessible at protocols.io: dx.doi.org/10.17504/protocols.io.eq2ly45dmlx9/v1 .No blinding or power analysis was required in this study.

## Ethics Statement

This article does not contain any studies with human or animal subjects performed by any of the authors.

