## Supplementary material for "Systematic Characterization of PBP2 as the Primary Siderophore Recognizer in Actinomycetes and Other Gram-Positive Bacteria": SI Appendix

Table S1. The information of 16 types of siderophore BGC

| Siderophore name | MIBiG accession[a] | SIDERITE ID[b] | Biosynthetic pathway | Genus | Reference |
|---|---|---|---|---|---|
| Bacillibactin | BGC0000309.5<br>BGC0002695.2<br>BGC0001185.5 | SID00152<br>SID00195<br>SID00151<br>SID00746<br>SID00747 | NRPS | *Bacillus* | [1, 2] |
| Petrobactin | BGC0000942.4 | SID00129 | NIS | *Bacillus* | [3] |
| Staphyloferrin A | BGC0000944.5 | SID00117 | NIS | *Staphylococcus* | [4] |
| Staphyloferrin B | BGC0000943.5 | SID00123<br>SID00125 | NIS | *Staphylococcus* | [5] |
| Heterobactin | BGC0002696.2;<br>BGC0000371.5 | SID00813<br>SID00594<br>SID00595<br>SID00597<br>SID00598; | NRPS | *Rhodococcus* | [6] |
| Dehydroxynocardamine | BGC0002073.3 | SID00018 | NIS | *Corynebacterium* | [7] |
| Desferrioxamine | BGC0002305.2<br>BGC0001478.5<br>BGC0001453.5<br>BGC0000940.5 | SID00012<br>SID00013<br>SID00016<br>SID00018<br>SID00020<br>SID00022<br>SID00027<br>SID00028<br>SID00030<br>SID00033<br>SID00335<br>SID00336<br>SID00338<br>SID00340<br>SID00344<br>SID00362<br>SID00655<br>SID00657<br>SID00772<br>SID00779<br>SID00727<br>SID00849 | NIS | *Streptomyces* | [8, 9] |
| Coelichelin | BGC0000325.5 | SID00305 | NRPS | *Streptomyces* | [10] |
| Scabichelin | BGC0000423.5 | SID00545 | NRPS | *Streptomyces* | [11] |
| Peucechelin | BGC0002466.3 | SID00298 | NRPS | *Streptomyces* | [12] |

| | | | | | |
|---|---|---|---|---|---|
| Fuscachelin | BGC0000359.5 | SID00222<br>SID00223<br>SID00596 | NRPS | *Streptomyces* | [13] |
| Albomycin delta2 | BGC0002300.2 | SID00560 | NRPS | *Streptomyces* | [14] |
| Griseobactin | BGC0000368.5 | SID00201 | NRPS | *Streptomyces* | [15] |
| Paenibactin | BGC0000401.5 | SID00196 | NRPS | *Streptomyces* | [16] |
| Qinichelins | BGC0001752.4 | SID00588<br>SID00589<br>SID00590<br>SID00591 | NRPS | *Streptomyces* | [17] |
| EDHA | BGC0002567.2<br>BGC0001587.5 | NA[c] | NIS-like[d] | *Streptomyces* | [18] |

a. In the MIBiG database, some siderophores have multiple accessions.

b. Certain siderophores exhibit structural variations due to different modifications, which correspond to multiple IDs in the SIDERT database.

c. This type of siderophore is not yet included in the SIDERT database.

d. This siderophore belongs to the aminopolycarboxylic acid type, and its biosynthetic pathway is NRP-independent; however, it does not contain canonical NIS synthetases.

Table S2.Number of SBPs in siderophore BGCs

| SBP | Number (128 unbiased siderophore BGCs) | Number (7,707 high-confidence siderophore BGCs) | Number (11,095 siderophore BGCs in total) |
|---|---|---|---|
| NMT1 | 0 | 3 | 12 |
| NMT1_3 | 0 | 0 | 33 |
| OpuAC | 6 | 128 | 148 |
| PBP_like_2 | 0 | 8 | 10 |
| Peripla_BP_1 | 0 | 8 | 13 |
| Peripla_BP_2 | 117 | 4626 | 5571 |
| Peripla_BP_3 | 3 | 93 | 161 |
| Peripla_BP_4 | 3 | 54 | 82 |
| Peripla_BP_5 | 0 | 2 | 2 |
| Peripla_BP_6 | 0 | 6 | 14 |
| SBP_bac_1 | 2 | 99 | 142 |
| SBP_bac_11 | 1 | 4 | 17 |
| SBP_bac_3 | 0 | 13 | 54 |
| SBP_bac_5 | 4 | 74 | 187 |
| SBP_bac_6 | 0 | 10 | 20 |
| SBP_bac_8 | 0 | 12 | 18 |
| TctC | 1 | 16 | 32 |
| ZnuA | 0 | 18 | 49 |

The table lists the counts of distinct SBPs across three dataset tiers: the curated unbiased dataset (n=128), the high-confidence dataset (similar to MIBiG, n=7,707), and the total dataset (n=11,095). Peripla_BP_2 consistently emerges as the predominant SBP across all datasets, exhibiting a frequency that is orders of magnitude higher than other common SBP such as SBP_bac_5, Peripla_BP_3, and OpuAC.

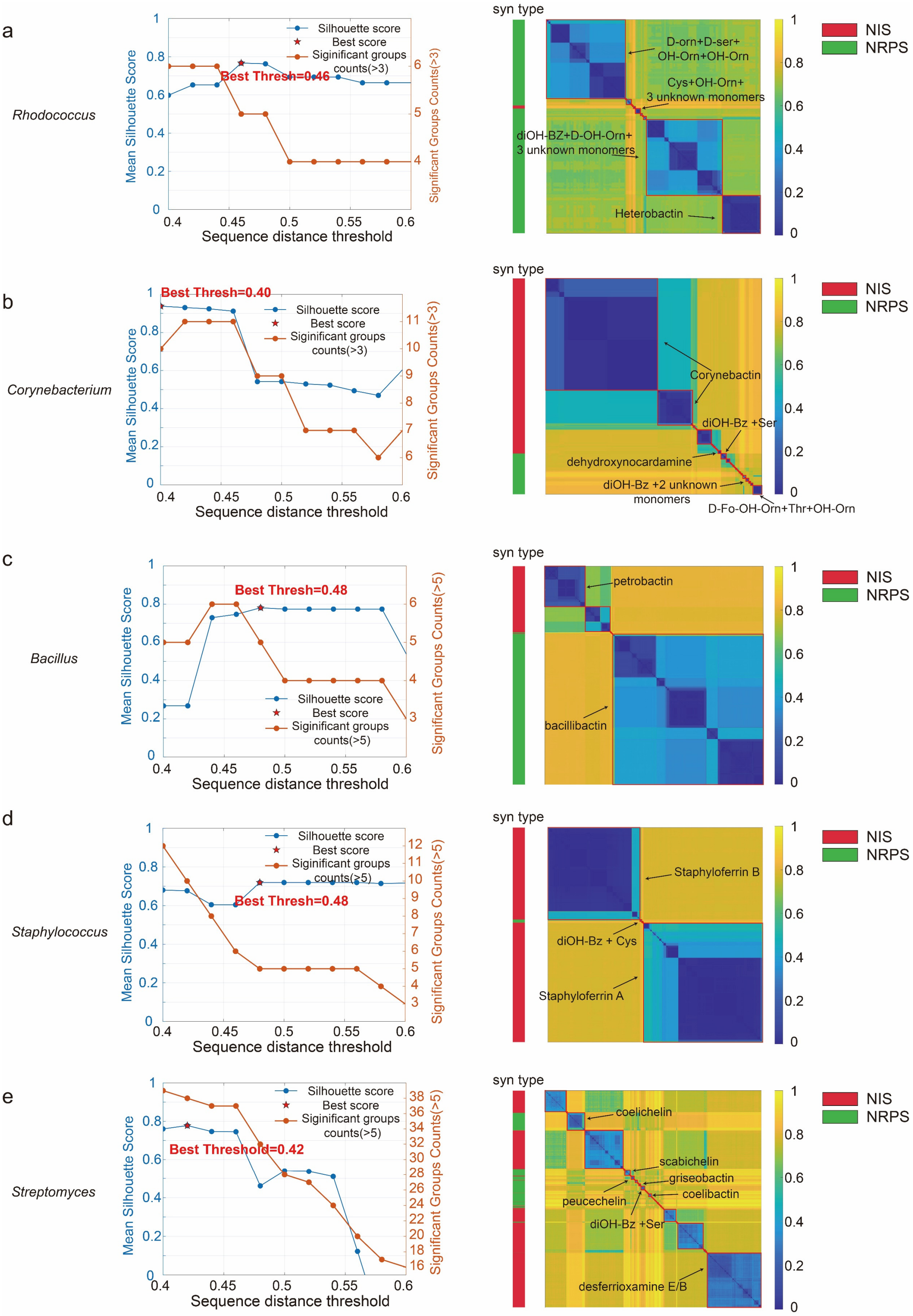


**Figure S1. Comprehensive clustering and characterization of siderophore BGCs.** **a-e** Clustering analysis for five representative genera: *Rhodococcus* (**a**), *Corynebacterium* (**b**), *Bacillus* (**c**), *Staphylococcus* (**d**), and *Streptomyces* (**e**). Left panels: Evaluation of optimal clustering thresholds. The line plots show the Mean Silhouette Score (blue) and the count of significant groups (orange) across varying sequence distance thresholds; the red star indicates the selected "Best Threshold" for each genus. Right panels: Heatmaps of pairwise distances between siderophore BGCs. Clusters are annotated with their predicted product or substrate monomers. The sidebar indicates the biosynthetic type (Red: NIS; Green: NRPS). Known siderophores (e.g., heterobactin, corynebactin, bacillibactin) are labeled, while novel clusters are annotated based on predicted monomer composition (e.g., "diOH-Bz + Ser")

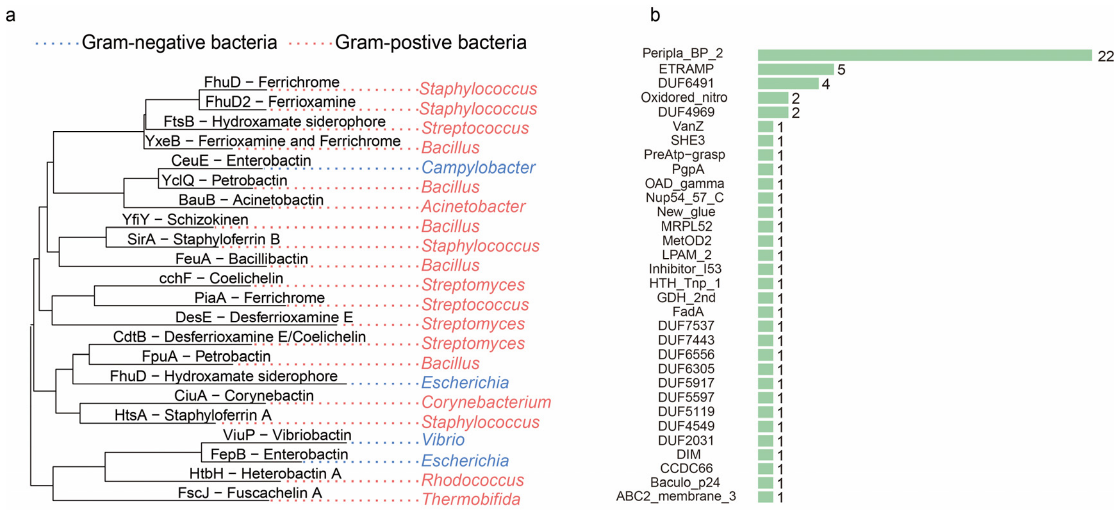


**Figure S2. Domain analysis of known siderophore cytoplasmic membrane recognizer.**

**a**. The phylogeny tree of 22 known siderophore recognizers.

**b**. The shared domain number among 22 known siderophore recognizers.

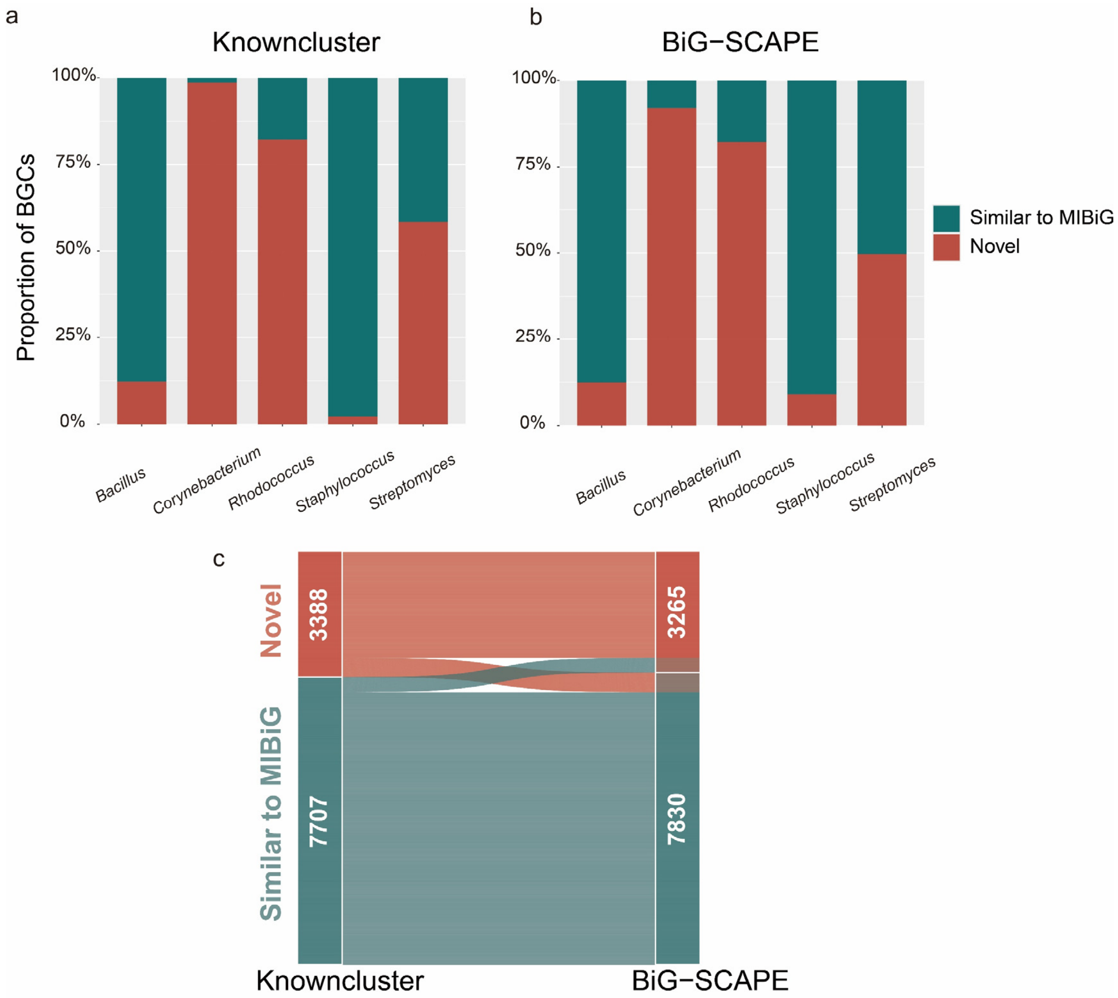


**Figure S3. Cross-validation of BGC novelty assessment using antiSMASH similarity versus BiG-SCAPE network analysis.**

**a**. Novelty classification based on the "Knowncluster" module (original method), where BGCs with >=80% similarity to a MIBiG entry are classified as "Similar to MIBiG" (teal), and those with < 80% are "Novel" (red).

**b**. Novelty classification based on BiG-SCAPE analysis (new method), where BGCs with a distance of <=0.3 to a MIBiG reference are classified as "Similar to MIBiG," and those with a distance > 0.3 are "Novel."

**c**. Alluvial diagram illustrating the high degree of concordance between the two methods. The flow connects the classification status of individual BGCs across the two approaches; 10,160 out of 11,095 BGCs (91.6%) retained the same classification, demonstrating that the novelty patterns are robust to different methodological thresholds.

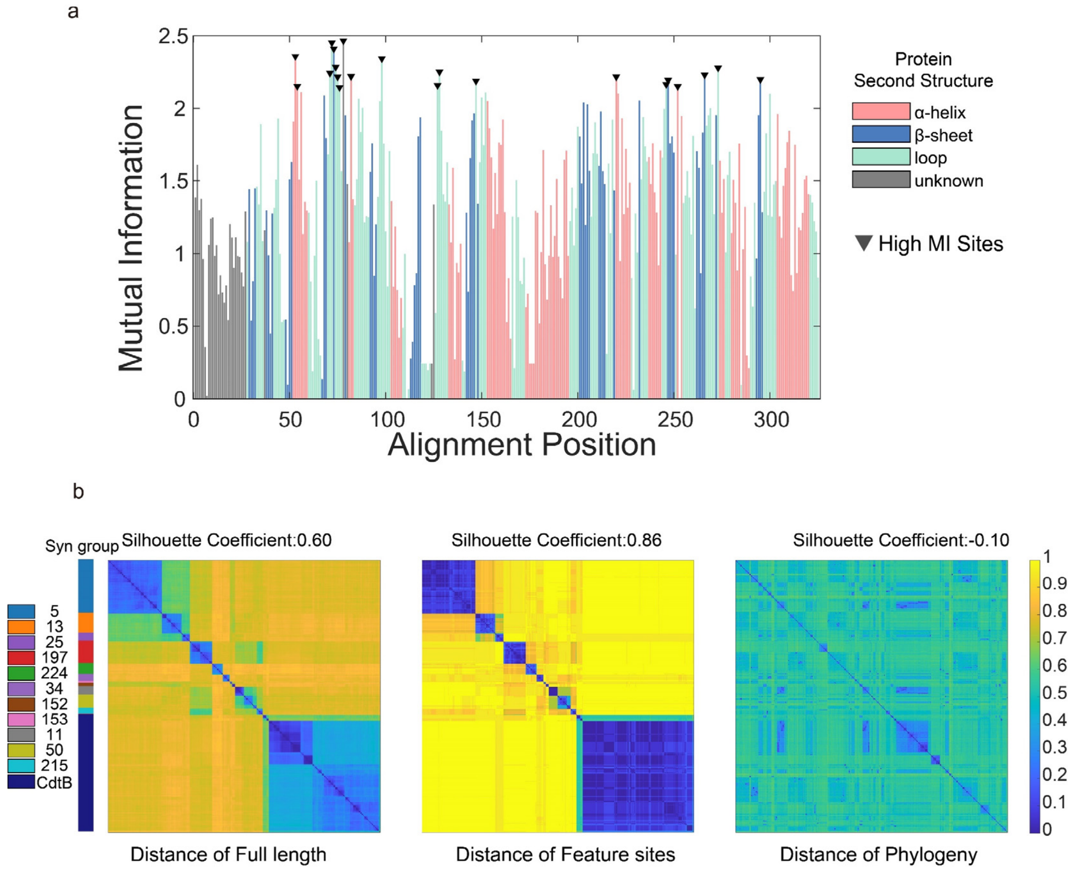


**Figure S4. Identification of feature sites of siderophore recognizers in *Streptomyces*.**

**a**. Mutual information analysis of 980 siderophore recognizer sequences from *Streptomyces*. Residues exhibiting mutual information values exceeding 80% of the theoretical maximum MI (reflecting strong correlation with recognizer grouping) are marked with black inverted triangles and defined as "feature sites."

**b**. Clustering performance comparison using different metrics. Pairwise distance heatmaps were generated for 980 *Streptomyces* PBP2 sequences using full-length sequence (left), feature sites (middle), and phylogenetic genes (right). The color bar indicates the sequence distance. The silhouette coefficient indicates the quality of clustering, with the feature-site-based metric achieving the highest score (0.86), demonstrating that the extracted feature sites effectively capture the variation responsible for functional specificity across 12 distinct siderophore recognizer groups. In the "Syn group" bar, groups are color-coded: numerical labels represent indices for distinct biosynthetic gene clusters (e.g., index 5 corresponds to the coelichelin BGC containing cchF, and index 224 corresponds to the desferrioxamine BGC), while "CdtB" denotes the experimentally characterized recognizer not colocalized with a siderophore BGC.

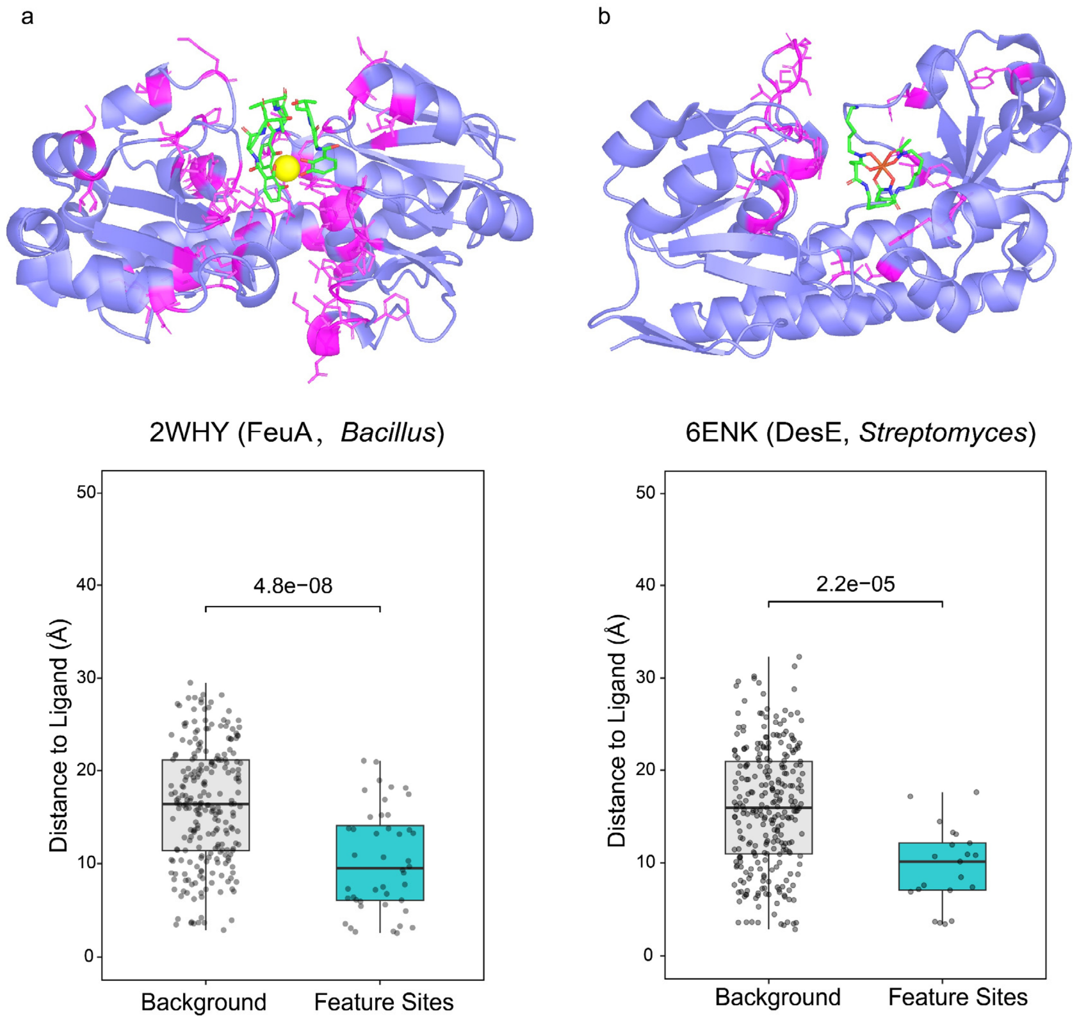


**Figure S5. Structural validation of specificity-determining sites in *Bacillus* and *Streptomyces*.**

**a,b.** Mapping of identified feature sites on representative crystal structures. Feature sites (magenta) identified by Mutual Information analysis are mapped onto the structure of the *Bacillus* recognizer FeuA (PDB: 2WHY) (left) and the *Streptomyces* recognizer DesE (PDB: 6ENK) (right). Note that in both genera, the high-MI sites structurally delineate the ligand-binding pocket. Boxplots quantifying the spatial distance between residues and the ligand. In both FeuA (left) and DesE (right), the identified feature sites are located significantly closer to the ligand compared to background residues. The *p* values were determined by a two-sided Wilcoxon rank-sum test

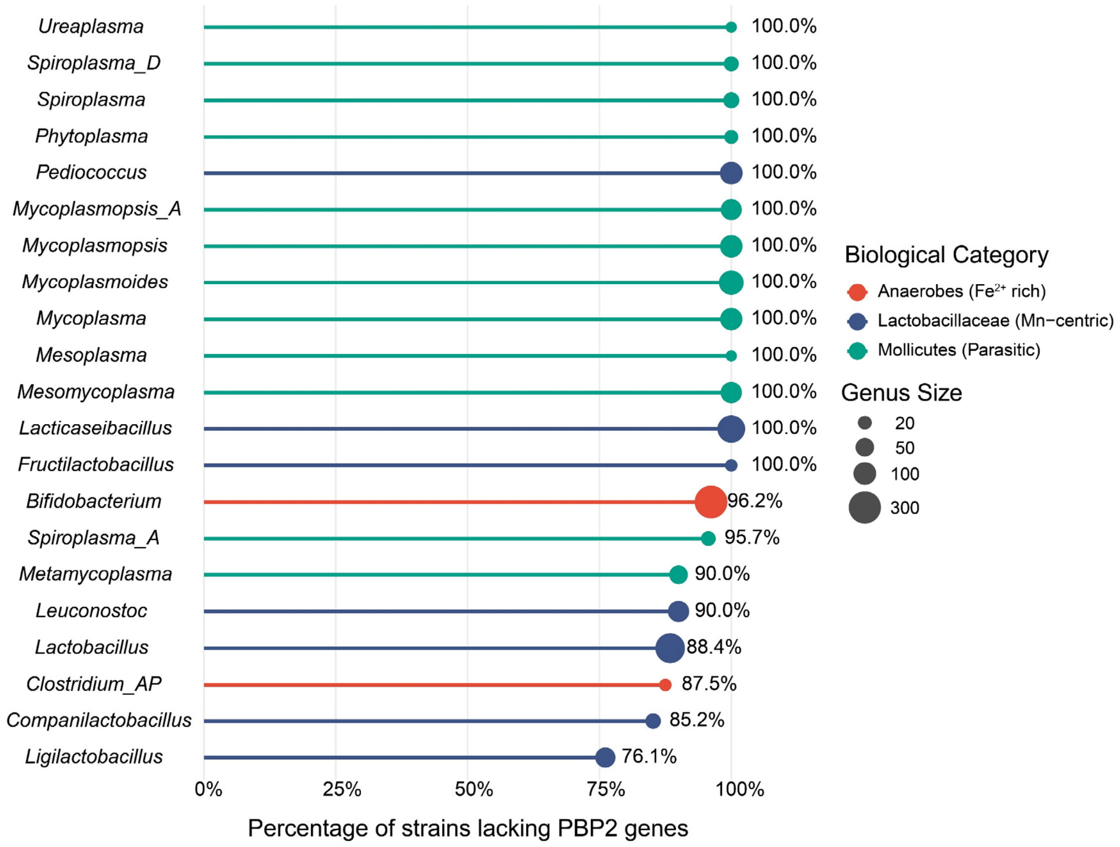


**Figure S6. Phylogenetic and ecological distribution of genera with PBP2 gene absence.**

The lollipop chart displays the percentage of strains lacking PBP2 genes. Genera are color-coded based on their physiological characteristics: Anaerobes (red, residing in $Fe^{2+}$-rich environments), lactic acid bacteria (blue, typically utilizing manganese as an iron substitute), and Mollicutes (green, parasitic lifestyle with reduced genomes). The size of each dot corresponds to the total number of genomes analyzed within that genus.

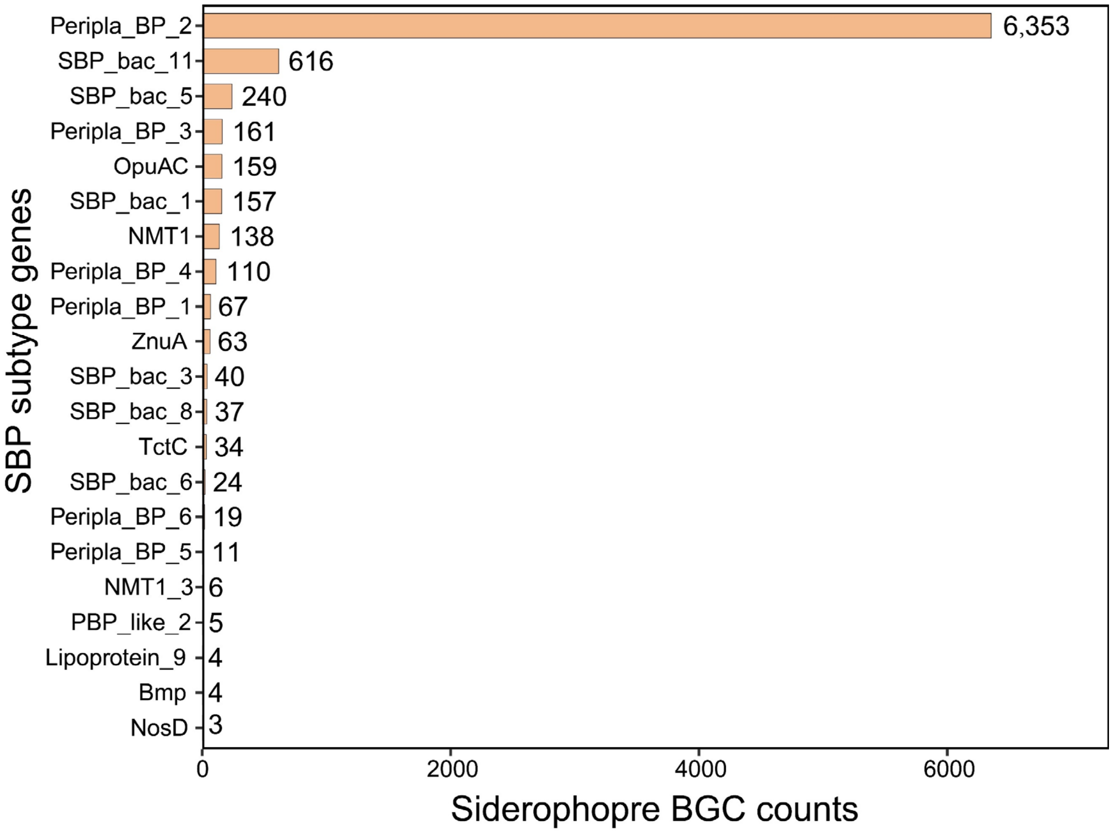


**Figure S7. Frequency of SBP genes within siderophore BGCs**.

The bar chart illustrates the abundance of different SBP genes identified inside 13,999 siderophore BGCs across the entire dataset of 16,232 Gram-positive genomes. The x-axis represents the total count of each SBP subtype found co-localized with siderophore BGCs, confirming PBP2 genes as the most prevalent SBP subtype genes associated with siderophore biosynthesis.

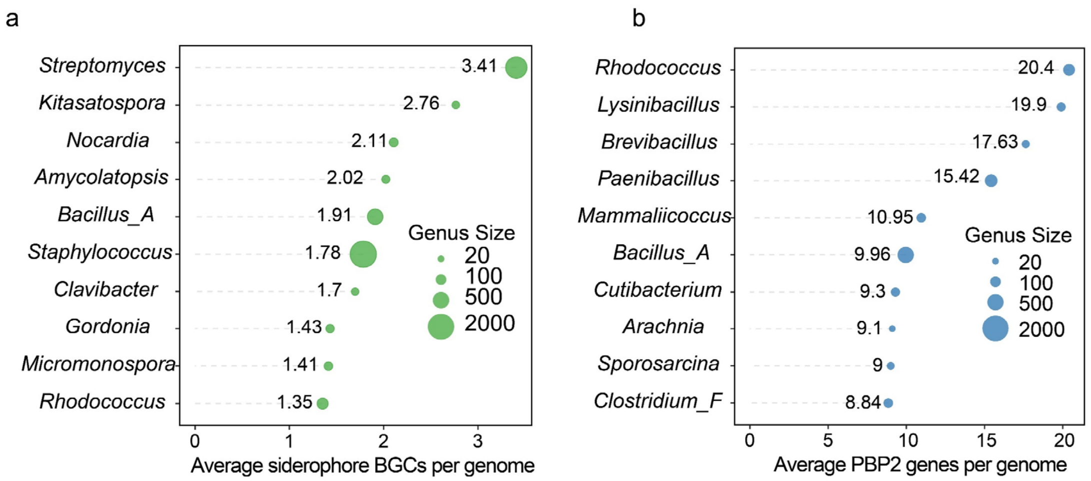


**Figure S8. Top genera ranked by siderophore biosynthetic and uptake potential.**

**a**. Top 10 genera ranked by the average number of siderophore BGCs per genome, representing high biosynthetic potential.

**b**. Top 10 genera ranked by the average number of PBP2 genes per genome, representing high uptake potential. In both panels, dot size corresponds to the number of genomes (genus size) for each genus.
